# Strategies to Identify and Edit Improvements in Synthetic Genome Segments Episomally

**DOI:** 10.1101/2022.03.18.484905

**Authors:** Alexandra Rudolph, Akos Nyerges, Anush Chiappino-Pepe, Matthieu Landon, Maximilien Baas-Thomas, George Church

**Affiliations:** Department of Genetics, Harvard Medical School, Boston, MA, 02115, USA; Wyss Institute for Biologically Inspired Engineering, Boston, MA, 02115, USA

## Abstract

Genome engineering projects often utilize bacterial artificial chromosomes (BACs) to carry multi-kilobase DNA segments on low copy number vectors. However, DNA synthesis and amplification have the potential to impose mutations on the contained DNA segments that can in turn reduce or prevent viability of the final strain. Here, we describe improvements to a multiplex automated genome engineering (MAGE) protocol to improve recombineering frequency and multiplexability. This protocol was applied to ‘recoding’ an *Escherichia coli* strain to swap out seven codons to synonymous alternatives genome-wide. Ten 44,402 to 47,179 bp *de novo* synthesized BAC-contained DNA segments from the recoded strain were unable to complement deletion of the corresponding 33 to 61 wild type genes using a single antibiotic resistance marker. Next-Generation Sequencing was used to identify 1-7 non-recoding mutations in essential genes per segment, and MAGE in turn proved a useful strategy to repair these mutations on the BAC-contained recoded segment when both the recoded and wild type copies of the mutated genes had to exist by necessity during the repair process. This strategy could be adapted to mutation identification and repair for other large-scale genome engineering projects, or for incorporation of small genetic engineering sites for quick protocol adjustments.

## INTRODUCTION

Since its description in the scientific literature over two decades ago (1), the use of single strand annealing proteins (SSAPs) to integrate short, single stranded DNA (ssDNA) oligonucleotides has undergone significant improvements to increase its efficiency as a genetic engineering tool. These include, but are not limited to, developing a protocol to simultaneously edit multiple sites using multiplex automated genome engineering (MAGE) (2), construction of broad-host plasmids with transient inactivation of mismatch repair (the pORTMAGE plasmid system) to enable higher efficiency recombineering across a range of host bacteria strains (3), and systematic identification of SSAPs that improve allelic replacement frequency in *E. coli* cells (4). Improvements to recombineering have been used for many applications, including, but not limited to, development of an *Escherichia coli* strain with all 321 instances of the UAG stop codon replaced genome-wide (5) and variant library production using directed evolution with genomic mutations (DIvERGE) to identify mutation combinations producing antibiotic resistance (6).

While recombineering with ssDNA oligonucleotides is a quick method to obtain an edited colony with low incidence of off-target mutations, improvements to recombineering efficiency prove important when the edit being introduced is deleterious, with unedited bacteria outcompeting edited bacteria. Additionally, the target location or desired edit may require synthesis of an oligonucleotide outside of optimal parameters, such as a particularly low folding energy, presence of a hairpin structure, or introduction of an insertion, deletion, or multiple edits on the same oligonucleotide that result in decreased homology of the oligonucleotide to the target genomic locus (2, 7, 8). Finally, certain applications may involve the presence of two or more loci with high homology to the oligonucleotide, such as the use of bacterial artificial chromosomes (BACs) in large-scale genome engineering work (9–11).

BAC recombineering is a well-established technique, especially for manipulating DNA used to develop transgenic animal or cell lines (12–15). Researchers have shown the technique’s capacity to incorporate double stranded DNA cassettes and ssDNA oligonucleotides (16–20). However, its utility in integrating ssDNA oligonucleotides for large-scale genome synthesis and engineering work, for which BACs provide a useful vector to hold and test long stretches of synthetic DNA, has not been well-established. Although many protocol parameters may be adjusted to improve recombineering efficiency, we tested the impact of four parameters on recombineering efficiency for a single recombineering cycle targeting the same locus: *E. coli* genomic background, oligonucleotide direction, SSAP selected, and cell count transformed.

The testing of these protocol parameters was applied to the repair of non-recoding mutations in *de novo* synthesized DNA segments used in the construction of a 57 codon recoded *E. coli* strain, *rE.coli*-57 (9). In genome recoding, all instances of one or more codons are replaced with synonymous alternatives in the genome and their corresponding translation machinery is removed to prevent recognition of the target codon (5, 9, 21). Successfully recoded organisms provide resistance to bacteriophage infection and horizontal gene transfer if the foreign DNA introduced contains the removed codons, and these organisms allow for strategic reintroduction of the removed codons with corresponding translation machinery encoding nonstandard amino acids for biocontainment and protein engineering purposes (9, 11, 22). When constructing the first recoded bacterial strain, C321dA, the 321 TAG stop codons in the *E. coli* MG1655 genome were recoded using MAGE in segment clusters and combined into a single strain using conjugative assembly genome engineering (CAGE) (5, 21). However, as researchers recode larger genomes or design more ambitious codon compression schemes, fully recoding a genome using MAGE becomes infeasible. Computational recoding, *de novo* DNA synthesis, and assembly on BACs to transfer into the genome becomes more practical (9, 10, 23).

In this work, we describe testing recombineering efficiency for cell count transformed, following testing to ensure selection of an optimal *E. coli* genotype, oligonucleotide direction, and pORTMAGE plasmid for introduction of a premature stop codon into the *lacZ* gene. Recombineering was applied to the development of a recoded *E. coli* strain with a seven codon compression scheme, *rE.coli*-57 (9). For this work, ten segments with a total of 22 non-recoding mutations in essential genes were selected as targets for repair on their BAC vectors using recombineering, allowing these segments to move to the strain assembly pipeline (9). Following repair of these non-recoding mutations, AlphaFold protein structure prediction was tested to determine whether the program could be used for hypothesis generation to address the question of why certain missense mutations improved complementation fitness.

## MATERIAL AND METHODS

### Bacterial Strains and Growth Conditions

Three *E. coli* strains were used to complete this project, selected because of their importance in the recombineering literature and the *rE.coli*-57 project: MDS42 (Scarab Genomics, full genotype can be found in Pósfai *et al.* 2006 (24)), TOP10 (Invitrogen, Cat. No. C404050, Genotype: F- *mcrA Δ(mrr-hsdRMS-mcrBC) Φ80lacZΔM15 ΔlacX74 recA1 araD139 Δ(araleu)7697 galU galK rpsL endA1 nupG StrR* (25)) and MG1655 (Genotype: F- lambda- *ilvG*- *rfb*-50 *rph*-1 (26)). Two of the three strains contained further adjustments to their genome. MDS42 has undergone CRISPR-mediated deletion of the *recA* gene to discourage unexpected homologous recombination. TOP10 has had its *lacZ* gene repaired using homologous recombination to allow for screening based on lactose fermentation on MacConkey agar with lactose. Each of these strains were grown overnight (12-18 hours) in Luria Broth-Lennox media with selective antibiotics (if applicable, for plasmid selection) at 32°C. For plating, overnight cultures were plated as dilutions on either Luria-Broth Lennox agar plates (if lactose fermentation is not being screened), or MacConkey agar with lactose plates (if lactose fermentation is being screened). Plates were incubated 1-6 days (depending on strain fitness) at 32°C.

### Oligonucleotides Used

A complete list of oligonucleotides used in this project is provided in the Supplementary Data (**Supplementary Table 1**). Kanamycin resistance cassette amplification primers and MASC primers were designed and described in a previous work (9). Primers were designed manually or using Geneious Prime 2022.0.2 Primer Design software to a Tm of 60-61°C. 90 bp MAGE oligonucleotides were designed to have a folding free energy between 0 and −15 kcal/mole, two phosphorothioated bonds on the 5’ ends, and have the desired mutation as centered as possible within the oligonucleotide, as per the literature (8). The *lacZ*-off oligonucleotides were designed to create a T35G mismatch in the *lacZ* gene, generating a V11* nonsense mutation featured in other recombineering projects (2, 4, 7).

### Plasmids Used

Synthesis and assembly of the 50 kb recoded segments from 2-4 kb Genebytes was described in a progress report on *rE.coli*-57 (9). These recoded segments were assembled on the pYES1L-URA BAC to allow for growth in both *Saccharomyces cerevisiae* and low copy number (1-2 copies per cell) maintenance in *E. coli*. The pYES1L-URA BAC is Spectinomycin-selectable in *E. coli*. Kanamycin deletions to test BAC-contained recoded segment complementation were performed using pKD78, a Chloramphenicol-selectable recombineering plasmid containing the three lambda-Red genes *exo*, *beta*, and *gam* activated through arabinose induction (9, 27)

Two recombineering plasmids were used during this project for ssDNA oligonucleotide recombineering. The Chloramphenicol-selectable pORTMAGE-4 recombineering plasmid contains the three lambda-Red genes and the dominant negative *mutL*E32K gene to inactivate the mismatch repair system transiently expressed using temperature induction (3). Finally, the Gentamycin-selectable pORTMAGE-503B recombineering plasmid contains the CspRecT SSAP and the dominant negative *mutL*E32K gene transiently expressed using m-toluic acid induction (4). Plasmids described are available through Addgene.

### Transformation Protocols

Transformation of electrocompetent bacteria was performed using protocols previously described in the literature (**Figure 1**). For transformation of recombineering plasmids, kanamycin resistance cassettes, and, for early recombineering experiments, 90 bp ssDNA oligonucleotides, a 1 mL transformation protocol described in Gallagher *et al.* 2014 was used (8). 3 mL of Luria Broth-Lennox media with selective antibiotics (if applicable, for the recombineering plasmid) was inoculated with the transformant strain and grown overnight (12-16 hours) at 32°C. The next day, a 1:100 dilution of this overnight culture was prepared in 3 mL Luria Broth-Lennox media with selective antibiotics (if applicable) and grown at 32°C to OD_600_ 0.3-0.5, as determined by spectrophotometry. If applicable, SSAP activation then proceeded, and the culture was chilled on ice for 20 minutes. 1 mL of the culture was then washed three times with chilled ultra-pure water, and, after the third wash, the cell pellet was resuspended in 80 μL of chilled ultra-pure water and 2-4 μL of the DNA to be transformed was added. 42 μL of the DNA:cell mixture was then added to a chilled 0.1 cm electrocuvette, and electroporated at 1.80 kV, 200 Ω, 25.0 μF. Cells were then recovered overnight in 1 mL Luria Broth-Lennox media and plated on Luria Broth-Lennox agar with selective antibiotics for 1-2 days at 32°C.

**Figure 1.**
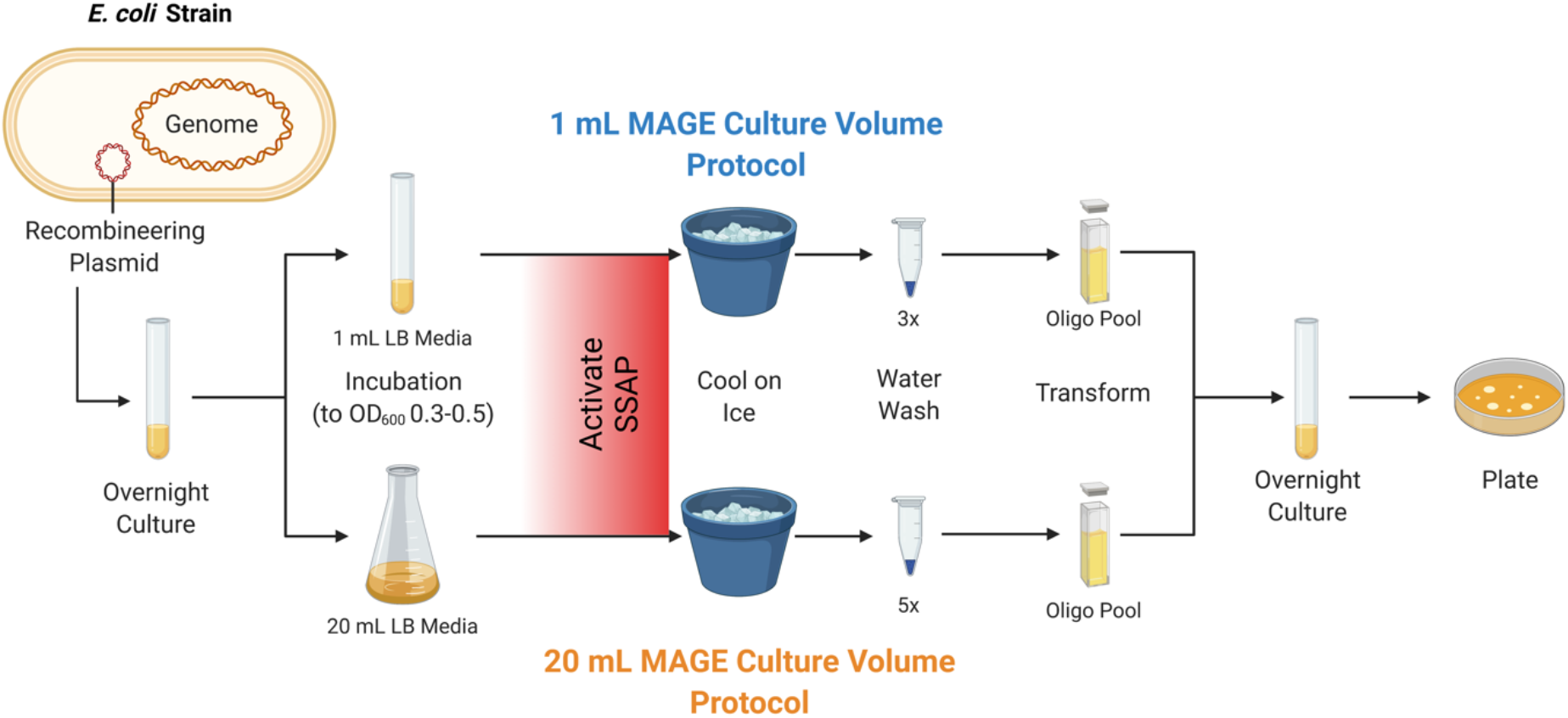
Overview of Recombineering Protocol. 90 bp oligonucleotides were recombineered into the strain of interest, an *E. coli* strain with a recombineering plasmid (either pORTMAGE-4 or pORTMAGE-503B) and, for certain applications, a 50 kb recoded segment on the pYES1L-URA BAC. Here, 1 mL (3 washes in water) or 20 mL (5 washes in water) MAGE protocols were used. Overnight recovery cultures were plated on Luria Broth-Lennox agar with antibiotic selection according to the pORTMAGE plasmid present, and repaired colonies were identified and quality checked with Sanger Sequencing and NGS, respectively. Figure was created with BioRender.com.

For transformation of 90 bp ssDNA oligonucleotides, a 20 mL transformation protocol described in Nyerges *et al.* 2016 was also used, with the increased cell pellet size found to be more researcher-friendly (3). 3 mL of Luria Broth-Lennox media with selective antibiotics for the recombineering plasmid was inoculated with the transformant strain and grown overnight at 32°C. The next day, a 1:100 dilution of this overnight culture was prepared in 25 mL Luria Broth-Lennox media with selective antibiotics and grown at 32°C to OD_600_ 0.3-0.5, as determined by spectrophotometry. SSAP activation then proceeded, and the culture was chilled on ice for 20 minutes. 20 mL of the culture was then pelleted and resuspended in 1 mL chilled ultra-pure water. This pellet was then washed five times with chilled ultra-pure water, and, after the fifth wash, the cell pellet was resuspended in 80 μL of chilled ultra-pure water and 2-4 μL of the DNA to be transformed was added. 42 μL of the DNA:cell mixture was then added to a chilled 0.1 cm electrocuvette, and electroporated at 1.80 kV, 200 Ω, 25.0 μF. Cells were then recovered overnight in 1 mL Luria Broth-Lennox media and plated on Luria Broth-Lennox agar with selective antibiotics for 1-2 days at 32°C.

A variation to resuspension of the final cell pellet was introduced to test for the impact of genomic background, oligonucleotide direction, SSAP selected, and cell count transformed. For testing the impact of genomic background, oligonucleotide direction, and SSAP selected, the 20 mL transformation protocol was used, and, following wash steps, the final cell pellet was instead resuspended in 500 μL water. For testing the impact of cell count transformed on recombineering efficiency, this 500 μL water resuspension volume was varied over a range of 100 μL to 1.3 mL. For both experiments, 240 μL of the cell:water resuspension was added to one or more new microcentrifuge tubes, and 12 μL of 500 μM oligonucleotide stock was added. Each 252 μL DNA:cell mixture was broken into five reactions of 42 μL into separate 0.1 cm electrocuvettes. Cell count transformed was calculated based on cell density per mL, outgrowth volume (20 mL), resuspension volume (intentionally varied over the range described), and volume of DNA:cell mixture added to each cuvette. Although cell division continues during activation, cell density was measured prior to SSAP activation, because of the importance of cell density measurements to determining when to activate the SSAP. Following recovery and plating, the total red and white colony count of each of these five reactions was combined as one replicate to increase the cells counted per reaction and to control for variability introduced by technical error on recombineering efficiency.

For colony sequencing, primers were manually generated to amplify the locus targeted for repair, and primers were designed to be allele specific to the recoded segment so that the corresponding genomic segment would not be sequenced. Polymerase chain reaction (PCR) was performed using 2GMP on the 30-192 post-recombineering colonies individually (average of 89 edited cells per sequencing run) and a wild type control (here, TOP10 or MDS42, for segment 12) to amplify the target locus and confirm allele specificity, respectively. Target DNA was sent to Genewiz (now Azenta Life Sciences) for Sanger Sequencing of unpurified PCR reactions.

For lactose fermentation screening, recovery cultures were plated on MacConkey agar containing lactose. Here, successful recombinants were unable to ferment lactose and resulted in white colonies (compared to lactose fermenters producing red colonies). Following recombineering and screening for white colonies on MacConkey agar media containing lactose, recombineering efficiencies were calculated by dividing the number of white colonies by the total number of colonies and multiplying by 100.

### Recoded Segment Complementation and Analysis

Complementation of corresponding wild type deletion by a BAC-contained recoded segment using a single kanamycin resistance cassette was tested using the methods and primers described in a previous work (9). A kanamycin resistance cassette with ~50 bp homology arms to the loci immediately flanking the corresponding wild type region was PCR amplified and gel purified. Then, an *E. coli* strain containing the recoded segment of interest on the pYES1L-URA BAC and the pKD78 recombineering plasmid was transformed with the purified kanamycin resistance cassette, recovered overnight, and plated on Luria Broth-Lennox agar containing kanamycin.

Determination of the result was based on the presence of colonies and the result from PCR testing. If colonies were not present for the whole segment deletion, the segment was diagnosed as requiring further troubleshooting to complement wild type deletion. If colonies were present, MASC-PCR was performed on colonies to check for presence of the corresponding wild type and recoded loci, as described in Ostrov *et al.* 2016 (9). If both wild type and recoded bands were fully present (eight bands indicating full presence of the corresponding locus), the segment was diagnosed as requiring further troubleshooting to complement wild type deletion (9).

### Next-Generation Sequencing (NGS) and Data Analysis

The recoded segment BACs were purified from an overnight 3-5 mL culture of the TOP10 *E. coli* strain containing the recoded segment on a pYES1L-URA BAC. BAC preps of the 10 segments described were sent to MiSeq for NGS to generate unpaired and paired 150 bp reads. Following read generation, read files were uploaded to Geneious Prime 2022.0.2 (http://www.geneious.com/). Reads were then processed using the Geneious “Trim and Filter” workflow, set to “Annotate new trimmed regions” with “Error Probability Limit” set to 0.05, and “Trim 5’ End” and “Trim 3’ End” selected. Filtered and trimmed reads were then aligned to recoded segment files using the Bowtie2 Geneious plug-in, local alignment setting (28).

### Computational Analysis of Protein Structure Predictions

Protein structure predictions were obtained using the DeepMind AlphaFold CoLab, running AlphaFold2.1.0. Protein amino acid sequences were input, and the notebook was set to “is_prokaryote” and “run_relax” settings. Runs were performed using a Google CoLab Pro+ account, with “High Ram Run” and “Run in Background” selected. Two protein alignments were generated for each wild type and mutated protein pair to generate the root mean square deviation (RMSD) for each (with higher RMSD values indicating more dissimilarity between the protein structures) and the template modelling score (TM-score) for RCSB (values ranging from 0 to 1, with 1 indicating the protein structures are identical). First, wild type and mutated protein versions were aligned using the RCSB Pairwise Structure Alignment tool, using the jFATCAT (rigid) algorithm. Next, wild type and mutated protein versions were aligned using the Pymol align command, running for five cycles to remove outlier atoms.

## RESULTS

### Testing Protocol Parameters Impacting Recombineering Efficiency

#### Testing the Impact of E. coli Genome, SSAP Selected, and Oligonucleotide Direction on Recombineering Efficiency

To determine the impact of strain characteristics on recombineering efficiency and to select characteristics resulting in the highest recombineering efficiency, different combinations of *E. coli* genomes, recombineering plasmids, and oligonucleotide directions were tested. Three common *E. coli* genomes used in genetic and genomic engineering work were used as the basis of strain construction: MDS42, TOP10, and MG1655 *E. coli* (24). For each of the three *E. coli* strains, two recombineering plasmids from the pORTMAGE system were incorporated via transformation: pORTMAGE-4 and pORTMAGE-503B. These plasmids were selected as each carried a different SSAP tested in the literature. While pORTMAGE-4 expresses the lambda-Red Beta SSAP, pORTMAGE-503B expresses the CspRecT SSAP (3, 4). The six strains constructed underwent a single cycle of recombineering to introduce a premature stop codon into the genomic *lacZ* gene, using either a forward or reverse direction version of the oligonucleotide to confirm the impact of oligonucleotide direction. Introduction of the premature stop codon via recombineering was screened for on MacConkey agar plates containing lactose.

Based on the scientific literature, we generated three hypotheses. First, the MG1655 *E. coli* strain will produce a higher recombineering efficiency than the TOP10 or MDS42 *E. coli* strains, as MG1655 is a more complete, wild type strain of *E. coli* and is commonly used for checking recombineering efficiency in *E. coli* (3, 4, 27, 29). Second, strains using the CspRecT SSAP will produce higher recombineering efficiencies than those using the Beta SSAP, as CspRecT is a high recombineering efficiency SSAP identified using a SSAP serial enrichment protocol (3, 4). Third, the reverse direction oligonucleotide will produce a higher recombineering efficiency than the forward direction oligonucleotide, as *lacZ* is on the first replichore of the *E. coli* genome on the negative strand (8).

Through characterization of the impact of *E. coli* genomic background, SSAP selected, and oligonucleotide direction, we were able to systematically confirm literature predictions for these factors. For all genomic background and SSAP combinations tested, the reverse direction oligonucleotide resulted in significantly higher recombineering efficiency than the forward direction oligonucleotide for genomic *lacZ* (**Figure 2A, 2B**). Beyond the oligonucleotide direction, the CspRecT SSAP resulted in higher recombineering efficiency for all *E. coli* strains when compared to their counterpart containing the Beta SSAP (**Figure 2B**). Overall, we saw that the MG1655 *E. coli* strain containing the CspRecT SSAP on the pORTMAGE-503B plasmid resulted in the highest recombineering efficiency when the oligonucleotide was properly designed as a reverse direction oligonucleotide to target the lagging strand for DNA replication (**Figure 2B**). Interestingly, a previous paper indicated that MDS42 outperformed TOP10 and MG1655 for plasmid uptake during transformation, indicating that the reduced recombineering efficiency seen here for MDS42 and TOP10 is specific to DNA incorporation into the genome (24).

**Figure 2.**
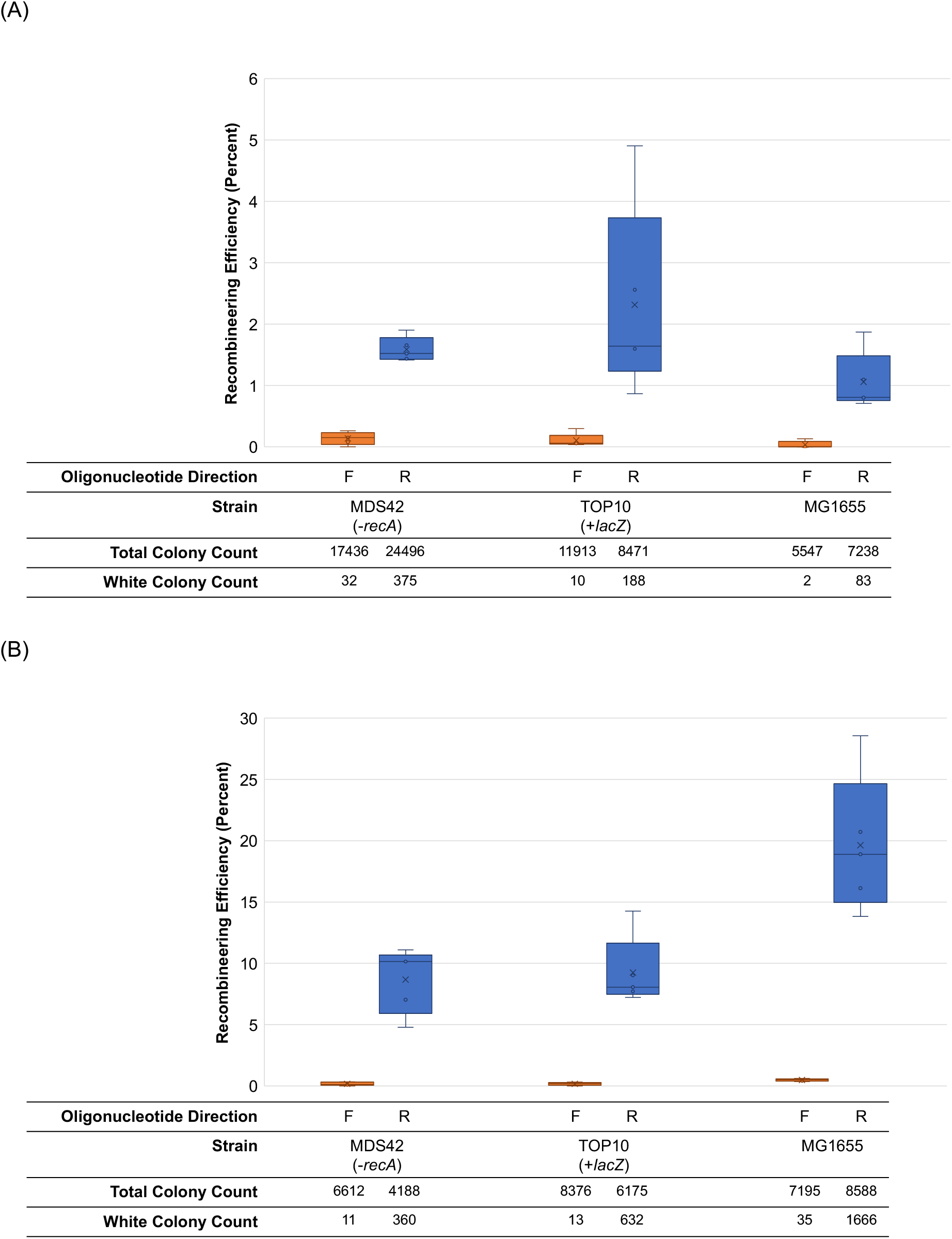
Impact of *E. coli* Genome, SSAP Selected, and Oligonucleotide Direction on Recombineering Efficiency. (A) One cycle of recombineering was performed to introduce a premature stop codon into the *lacZ* gene, recovered overnight in Luria Broth-Lennox media, and plated on MacConkey agar with lactose. Recombineering was performed targeting the genomic copy of *lacZ* in three *E. coli* genomes: MDS42(-*recA*), TOP10 (with *lacZ* repaired), and MG1655. An identical oligonucleotide was developed in the forward and reverse directions, and the pORTMAGE-4 plasmid was used to supply the Beta SSAP. (B) The previous experiment was repeated, using the pORTMAGE-503B plasmid to supply the CspRecT SSAP. Orange boxes correspond to forward (F) oligonucleotide direction, while blue boxes correspond to reverse (R) oligonucleotide direction.

#### Testing the Impact of Cell Count Transformed on Recombineering Efficiency

As the MG1655 *E. coli* strain, CspRecT SSAP, and reverse direction oligonucleotide combination resulted in the highest recombineering efficiency when targeting the genomic *lacZ* gene, these parameters were then used to test the impact of cell count transformed on recombineering efficiency (**Figure 1**). By testing a range from 1.67*10^8^ cells per cuvette to 3.07*10^9^ cells per cuvette, we found that between 3.20*10^8^ cells per cuvette and 8.32*10^8^ cells per cuvette results in the highest recombineering efficiency (**Figure 3A, 3B**).

**Figure 3.**
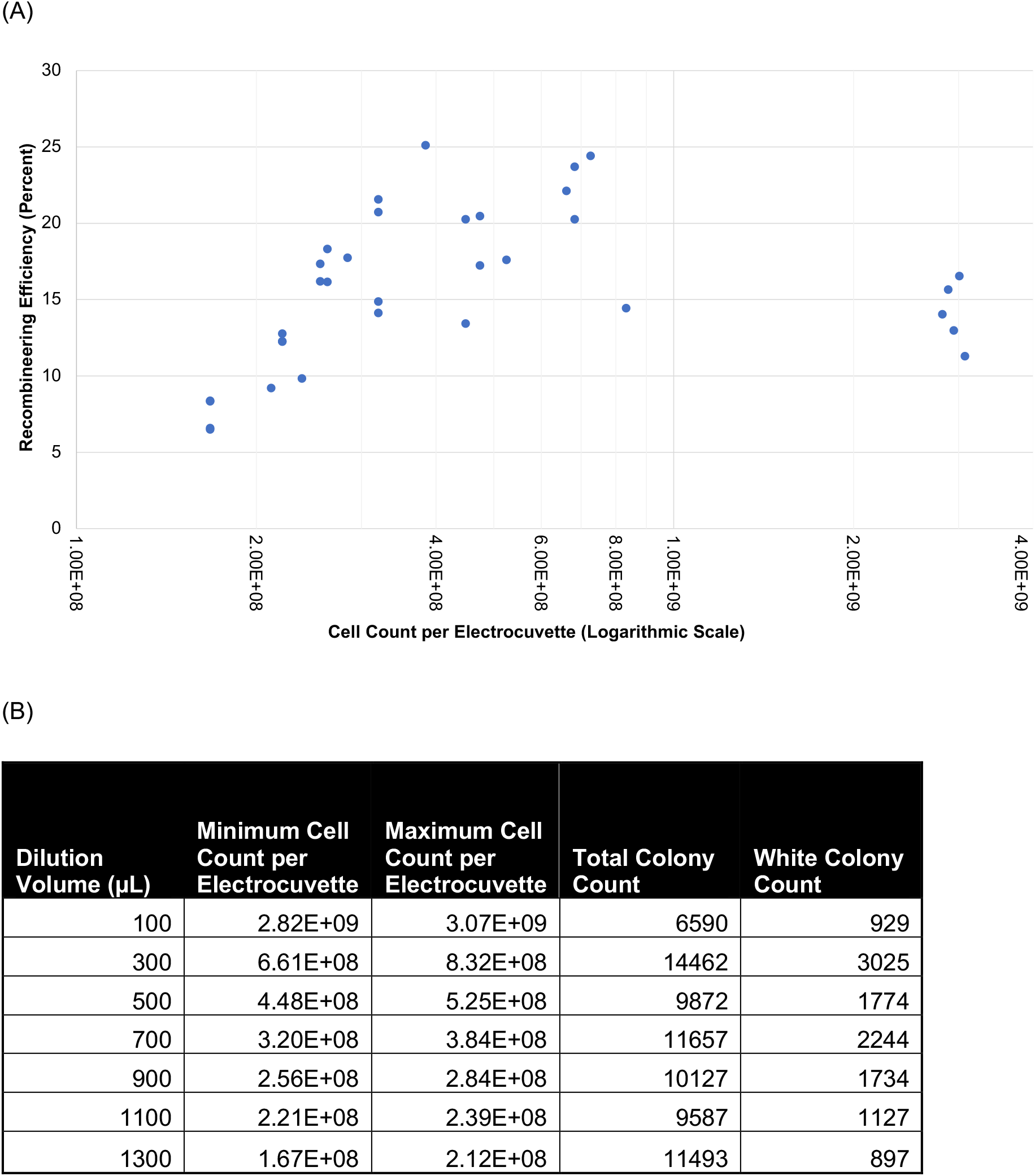
Impact of Cell Count Transformed on Recombineering Efficiency. (A) MG1655 *E. coli* with pORTMAGE-503B underwent one cycle of recombineering using the reverse direction oligonucleotide introducing the premature stop codon into the genomic *lacZ* gene. Different cell counts were transformed to determine the optimal cell count for transformation per electrocuvette, achieved by diluting the final cell pellet after washing in different volumes of ultra-pure water. (B) Values for minimum and maximum cell count per electrocuvette, and total and white colony count associated with each dilution volume on MacConkey agar plates with lactose.

Cells per cuvette was chosen as a parameter to test, because increasing the amount of water the final cell pellet is diluted in significantly changes the number of possible transformations from a single washed culture, assuming 40 μL of the final cell resuspension in water is used consistently for transformation with 1-2 μL 500 uM oligonucleotide. As between 3.20*10^8^ cells per cuvette and 8.32*10^8^ cells per cuvette corresponds to a 20 mL culture MAGE protocol with the final cell pellet diluted in 300 to 700 μL water, this result demonstrates that up to 17 parallelized recombineering reactions can be performed simultaneously to obtain a high recombineering efficiency with a minimal increase in effort compared to that needed for one recombineering reaction, as only the steps following the MAGE cell pellet wash expand to accommodate multiple transformations. Furthermore, while increasing the amount of water the final cell pellet is diluted in beyond 700 μL does lower the recombineering efficiency, further dilution increases the number of parallelized recombineering reactions possible for a single 20 mL culture. For dilution in 900 μL water, recombineering efficiency is at an average of 17.2%, with a 900 μL dilution allowing for 22 parallelized recombineering reactions. Further dilution may even be possible for beneficial mutations, such as those resulting in overcoming antibiotic selection for growth on media.

### BAC Recombineering to Repair Mutations in Episomal *de novo* Synthesized DNA Segments

#### Requirement for BAC Recombineering in rE.coli-57 Strain Construction

Rigorous optimization of the recombineering protocol has many possible applications, with an important one being the construction of synthetic genomes. Here, we apply recombineering to editing BACs containing MDS42 *E. coli* genome segments towards the development of a recoded *E. coli* strain (*rE.coli*-57). The *rE.coli*-57 genome was designed computationally to optimize synonymous replacement of 62,214 instances of the seven codons from the MDS42 *E. coli* genome, to allow for later deletion of their five corresponding tRNAs and one release factor (9). To ease troubleshooting, the computationally designed genome was then divided into 87 segments, approximately 40 genes (or 50 kilobases) long constructed from 2-4 kb overlapping Genebytes on BACs with a mini-F replication origin (pYES1L- URA) (9). To determine whether recoded segments were able to complement wild type deletion, recombineering-mediated deletion with a single antibiotic resistance cassette was performed to remove the corresponding wild type genes in the TOP10 host genome (9). For recoded segments unable to complement wild type deletion, NGS of the BAC-contained, synthesized segment was used to identify <30 bp non-recoding mutations present in recoded essential genes (30). In this manner, ten recoded segments were identified as candidates for non-recoding mutation repair (summary in **Table 1**, full list of mutations identified in **Supplementary Table 2**). As the recoded segment was contained on a BAC for transfer from *S. cerevisiae* into *E. coli*, BAC recombineering was identified as the strategy to use for non-recoding mutation repair (9). This BAC recombineering would be performed on recoded segment copies contained on BACs in a TOP10 *E. coli* strain, with a minimum of two target copies present per strain as the wild type copy of the segment could not be deleted prior to recombineering, thus necessitating higher efficiency recombineering despite the need to obtain only one strain with the recoded segment successfully repaired.

**Table 1.**
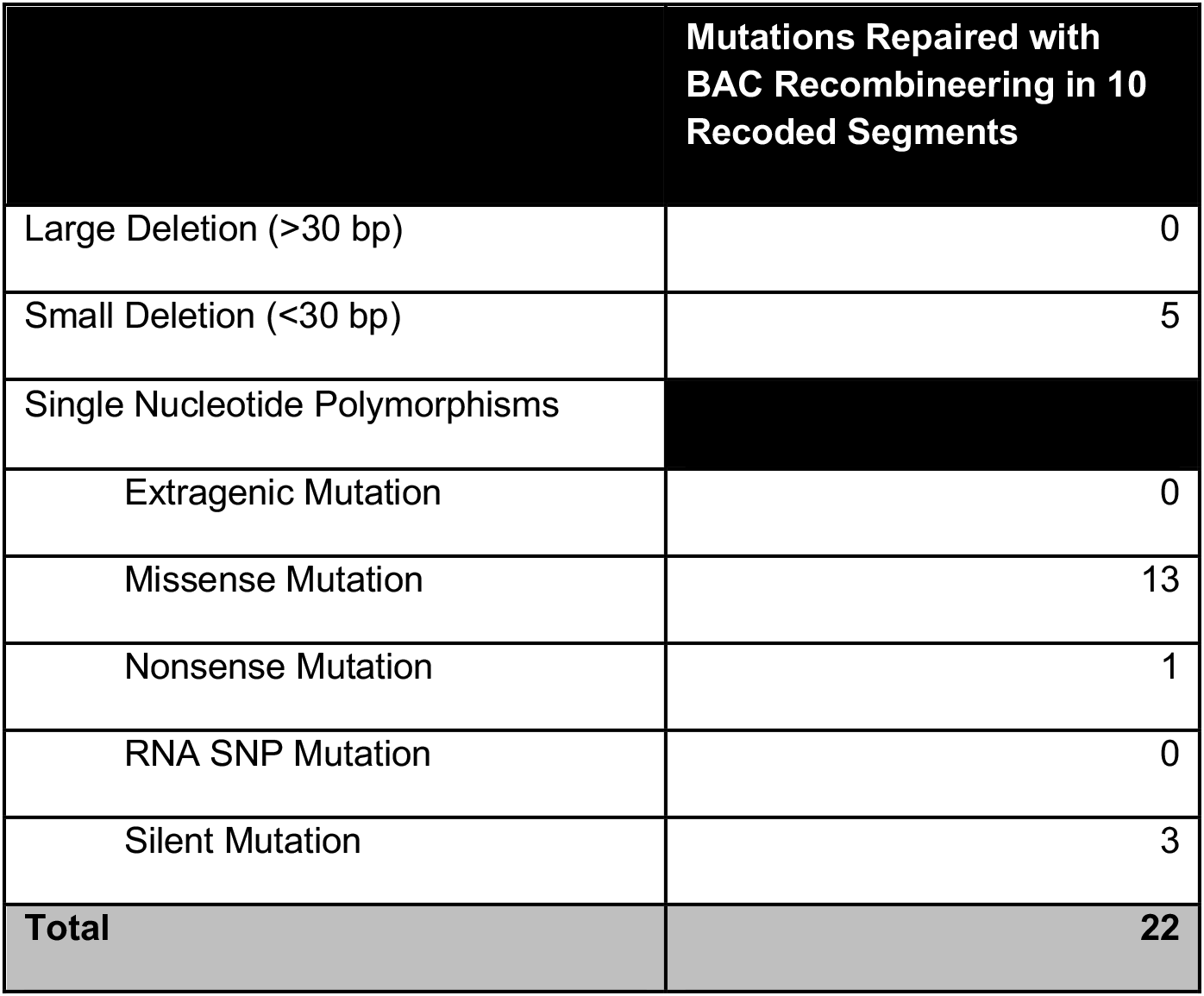
Overview of non-recoding mutations present in ten *rE.coli*-57 segments repaired using BAC recombineering.

#### Non-Recoding Mutation Repair Strategy

BAC recombineering was used to repair single nucleotide polymorphisms (SNPs) and small deletions ranging from 1-30 bp in ten synthesized, recoded segments using the pORTMAGE recombineering plasmid system. For nine segments, non-recoding mutations present in recoded essential genes were targeted for repair on a segment copy contained on a BAC in a TOP10 *E. coli* host, as TOP10 is an *E. coli* strain suited to maintaining clonal DNA. For one segment (segment 12), repair was instead performed on the same BAC in an MDS42(-*recA*) strain, as the TOP10 genomic locus containing the segment genes overlapped with a large repeat region not present in the MDS42 strain.

#### Single Oligonucleotide BAC Recombineering to Repair a dnaG Missense Mutation

One of the recoded segments repaired was recoded segment 59. While segment 59 contains four essential genes (*rpsU*, *dnaG*, *rpoD*, and *higA*), one essential gene (*dnaG*) was found to contain a non-recoding mutation based on NGS (**Figure 4A**). The P470L missense mutation in the DnaG primase protein is located in the C-terminal domain of the protein, specifically within the hydrophobic pocket that interacts with the C-terminal tail of the SSB protein (31).

**Figure 4.**
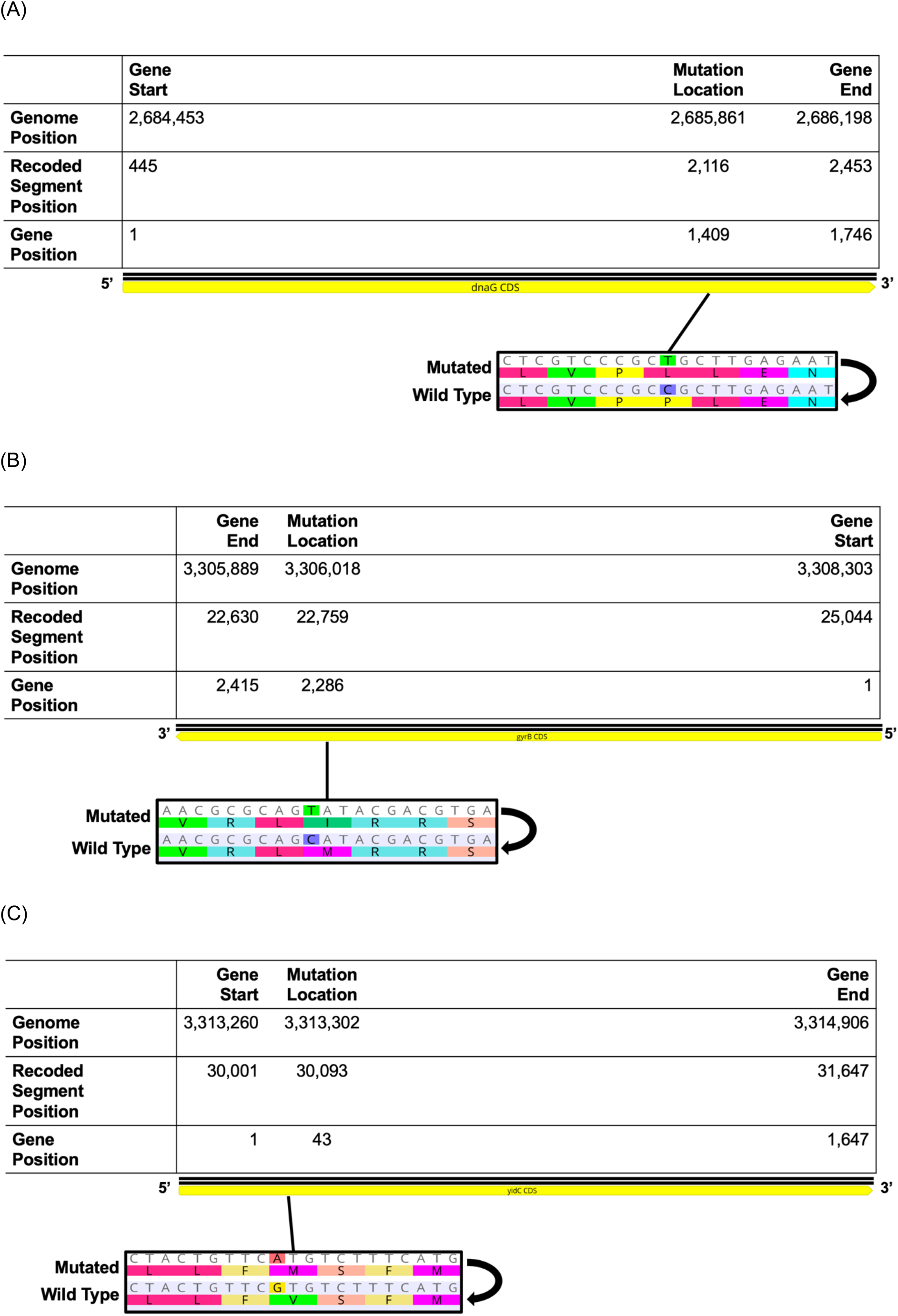
NGS Identifies SNPs in Essential Genes in Recoded Segments 59 and 72. (A) The recoded *dnaG* gene in segment 59 contains a C to T SNP, resulting in a P470L missense mutation in the DnaG protein product. (B) The recoded *gyrB* gene in segment 72 contains a C to T SNP, resulting in a M762I missense mutation in the GyrB protein product. (C) The recoded *yidC* gene in segment 72 contains a G to A SNP, resulting in a V15M missense mutation in the YidC protein product. Genome and segment positions are given based on the published *rE.coli-*57 genome (9). Figure was created with Geneious Prime 2022.0.2.

The mutation was repaired on the BAC-contained recoded segment 59 using one cycle of MAGE in a TOP10 strain with pORTMAGE-4 as the recombineering plasmid, with a recombineering efficiency of 3.2%. Following recombineering-mediated repair, Sanger Sequencing was used to obtain a copy of the recoded segment with the missense mutation repaired in *dnaG*, and NGS was used to confirm no additional missense mutations were present in recoded essential genes following repair. Complementation by the whole recoded segment was then tested, and it was found that the repaired segment 59 could now complement wild type deletion.

#### Multiple Oligonucleotide BAC Recombineering to Repair gyrB and yidC Missense Mutations

While the repair of *dnaG* in recoded segment 59 served as an example where only one missense mutation was present in a recoded essential gene in a segment, the average number of mutations repaired with BAC recombineering per 44,402 to 47,179 bp segment was 2.2 mutations. Recoded segment 72 was a synthesized DNA segment for which multiple non-recoding mutations were repaired simultaneously, here two repairs. While segment 72 contains six essential genes (*gyrB*, *dnaN*, *dnaA*, *rpmH*, *rnpA*, and *yidC*), only two were found to contain non-recoding mutations: *gyrB* and *yidC* (**Figure 4B, 4C**). The M762I missense mutation in GyrB was in the C-terminus of the protein, outside of catalytic domains. Meanwhile, the V15M missense mutation in YidC was located in the N-terminus of the protein, in the first transmembrane domain signal-anchor sequence (32).

The *gyrB* and *yidC* missense mutations were repaired simultaneously with an oligonucleotide pool on the BAC-contained recoded segment 72. Four cycles of MAGE were performed in a TOP10 strain with pORTMAGE-503B as the recombineering plasmid, with recombineering efficiencies of 23.4% for *gyrB* and 28.1% for *yidC*. Repair of the mutation was confirmed as described for segment 59. However, although recoded segment 72 could now complement wild type deletion, it did so with decreased fitness, indicating that other avenues for segment troubleshooting remain to be explored. We hypothesize that troubleshooting of the recoding scheme for the segment will lead to further fitness improvements for segment 72 complementation. By performing BAC recombineering on recoded segment 72 with the wild type copy of the segment deleted to test candidates for improvement, later research can take advantage of the presence of only the recoded segment for improvements.

### Assessing Use of Protein Structure Prediction to Prioritize Non-Recoding Mutation Repair with Test Segments

Following completion of the repair of 22 non-recoding mutations in ten recoded segments’ essential genes, we were interested in whether currently available computational programs could allow future efforts for large-scale genome engineering to prioritize repair of unexpected mutations, particularly those resulting in missense mutations. While working on this project, the DeepMind team released AlphaFold, allowing for robust protein structure predictions from amino acid sequences (33, 34). Although this enormous undertaking represents a significant leap forward in computational biology, recent work has indicated the need for cautious optimism regarding the use of AlphaFold to predict the impact of individual mutations on a protein, encouraging researchers to bear in mind that protein function and protein structure are not equivalent (35, 36).

Here, we look at predictions made regarding protein structure on the two example segments described above. In the first example, repair of a single missense mutation in segment 59’s *dnaG* improved complementation fitness. However, while repairing two missense mutations in recoded segment 72 simultaneously improved fitness, the impact of these individual mutations on recoded segment complementation fitness is unknown. Here, we tested the use of AlphaFold to predict whether it was the repair of one or both mutations that improved complementation fitness. We used the AlphaFold CoLab notebook for these predictions to encourage researchers with all levels of computational background to use such tools for their own projects.

Wild type and mutated protein structure predictions were aligned using RCSB Pairwise Structure Alignment (using a jFATCAT-rigid alignment) and Pymol, and protein structure similarity was assessed using RMSD values (37–40) (**Figure 5**). Importantly, while RCSB Pairwise Structure Alignment reports RMSD without removing outlier atoms, Pymol align reports RMSD both before and after a set number of cycles filtering for outlier atoms. Here, we report the RCSB Pairwise Structure Alignment RMSD, as well as both the Pymol RMSD values before outlier filtering and after five cycles of outlier filtering (**Table 2**). However, all RMSD values generated were very low, indicating the protein structures compared in the alignments were very similar. This was expected due to the high degree of protein sequence homology between mutated and wild type protein sequences. RMSD assessed with all atoms (RCSB RMSD and PyMOL RMSD without refinement cycles) indicated DnaG and YidC wild type and mutated proteins were more dissimilar, while GyrB’s wild type and mutated versions were more similar. However, by applying five refinement cycles for RMSD calculation with PyMOL, the conclusion changes to GyrB and YidC wild type and mutated versions being more comparable, while DnaG’s mutated and wild type forms were more dissimilar. Therefore, RMSD does not allow us to form a hypothesis about which of these amino acid mutations has the greatest impact on protein structure, and thus is the likely candidate for repair.

**Table 2:** RMSD and TM-Score for DnaG, YidC, and GyrB Wild Type and Mutated Predicted Protein Structure Alignments.

| Protein Name | Protein Length (aa) | RCSB Protein Alignment RMSD (Angstroms) | PyMOL Protein Alignment RMSD (Angstroms) |  | TM-Score |
| --- | --- | --- | --- | --- | --- |
|  |  |  | 0 Cycles | 5 Cycles |  |
| DnaG | 581 | 0.75 | 0.75 | 0.63 | 0.99 |
| GyrB | 804 | 0.35 | 0.35 | 0.23 | 1 |
| YidC | 548 | 0.66 | 0.66 | 0.24 | 0.99 |

**Figure 5.**
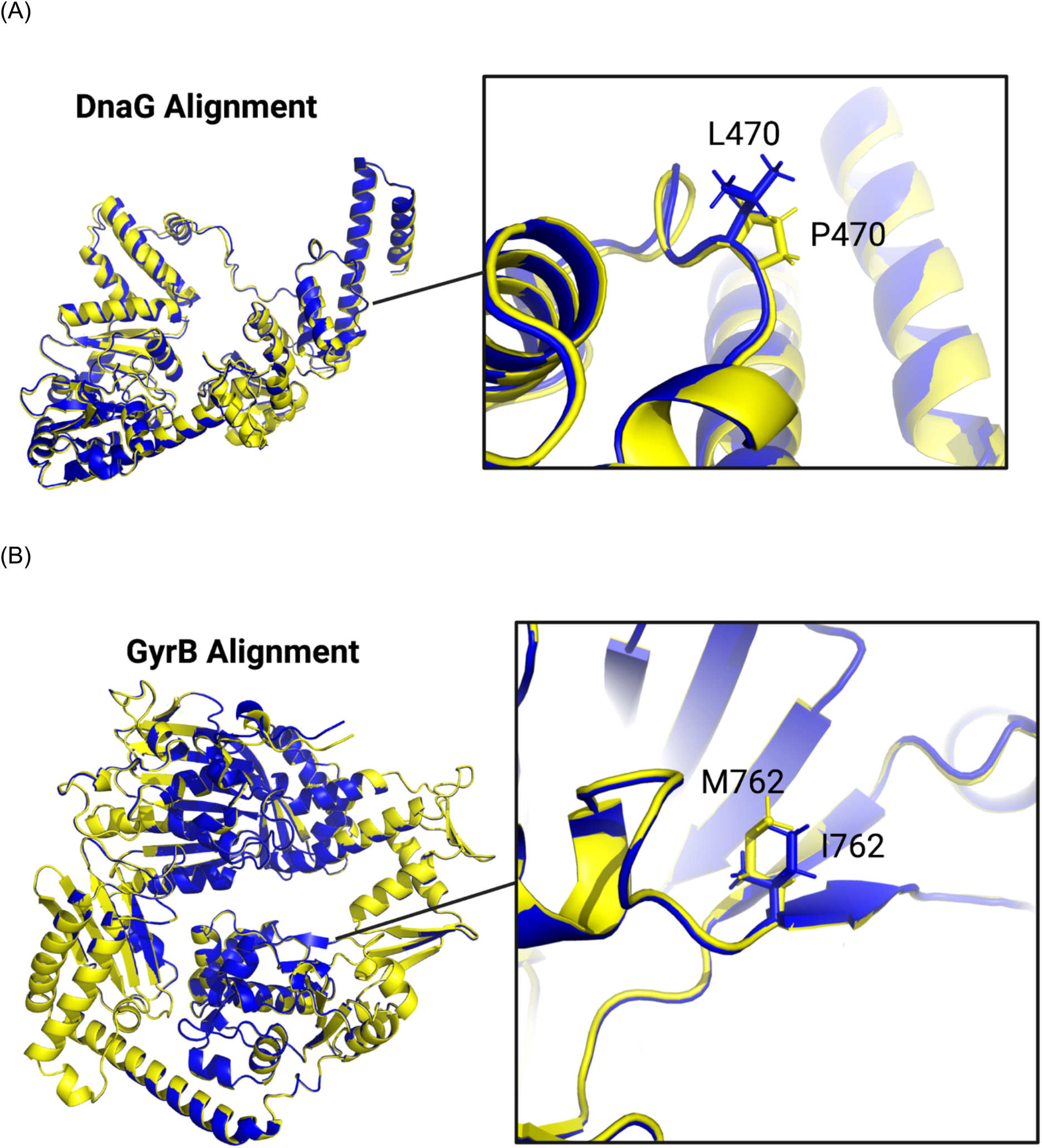
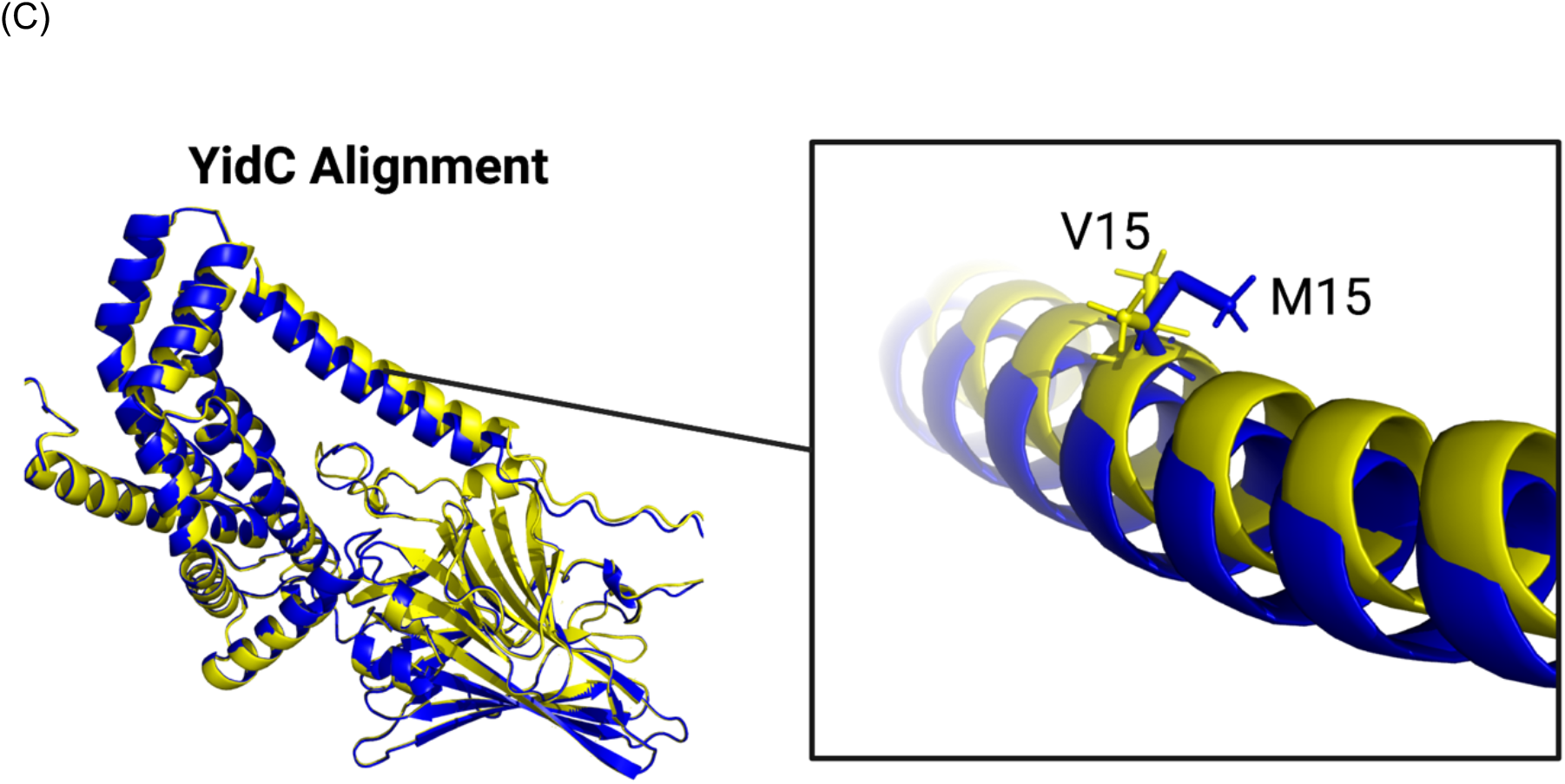
Protein Structure Prediction Alignments for Wild Type and Mutated DnaG, GyrB, and YidC. Protein structure prediction PDB files obtained from the AlphaFold2 CoLab Notebook were aligned using PyMOL. Wild type structures are shown in yellow, and mutated structures are shown in blue. Wild type and mutated residues are shown as insets, with zoom set to 12 angstroms. (A) DnaG wild type (P470) and mutated (L470) protein structure alignment. (B) GyrB wild type (M762) and mutated (I762) protein structure alignment. (C) YidC wild type (V15) and mutated (M15) protein structure alignment. Figure was created with PyMol.

As the RMSD values are impacted by sequence length, TM-score was also used to determine whether sequence length impacted our ability to use protein structure predictions to test the impact of single residue changes on protein structure (41). However, we again saw that the TM-scores for all protein alignments between wild type and mutated forms were very similar, with the alignment for GyrB having a TM-score of 1 and the alignments for DnaG and YidC both having TM-scores of 0.99. In this specific test case, this tool does not allow us to generate a hypothesis as to which mutation should have been prioritized for repair in recoded segment 72. However, as researchers seek to answer different questions, this result should not be intended to discourage other teams from using AlphaFold for their own test cases or to see if the problem posed in this section can be addressed with a larger dataset.

## DISCUSSION

Here, we tested four parameters of the recombineering protocol to increase recombineering efficiency for ssDNA oligonucleotide incorporation into the *E. coli* genome. This optimization was applied to repairing 22 non-recoding mutations in ten 44,402 to 47,179 bp recoded segments contained on BACs towards completion of a recoded *E. coli* strain with a seven-codon compression scheme. As anticipated when starting this project, repair of non-recoding mutations improved fitness for some, but not all, recoded segments described, thus indicating that other problems may contribute to reduced segment fitness. For this work, we repaired non-recoding mutations in essential genes, but acknowledge that different methods of identifying essential genes result in some variation in the list of genes defined as essential, and gene essentiality cannot be categorized in a binary manner (42–44). Additionally, for many applications of complementation testing for synthetic DNA segments against wild type counterparts, complementation by the synthetic DNA segment may not entirely eliminate fitness, but reduce fitness compared to wild type. While the segments of interest in this paper were those for which complementation by the recoded segment caused fitness loss, it would be possible to use a similar strategy to the one described in this paper to repair segments for which complementation reduced, but did not eliminate, fitness. Quantification of fitness using a plate reader growth assay would be optimal to determine the extent to which repair efforts improve fitness of a strain forced to rely on the synthetic segment compared to its wild type counterpart. Identification and repair of further causes of decreased complementation fitness is an exciting challenge, with the possibility of uncovering novel biological rules.

Further, as this work was done in the context of constructing a fully recoded *E. coli* strain, the non-recoding mutations identified and repaired were not the only changes made to the MDS42 genome. Previous recoding efforts have identified possible means by which recoding may require further troubleshooting, including changes to mRNA folding and ribosomal binding sites (10, 45). While identification of individual recoded codons impacting fitness is a non-trivial task, the BAC recombineering strategy described in this work could be used to repair both recoding and non-recoding mutations contributing to decreased fitness. Additionally, repair of non-recoding mutations generated randomly during *de novo* DNA synthesis allows researchers to opportunistically identify novel mutations impacting strain fitness that can, in turn, direct hypothesis generation and testing. By performing large-scale genome engineering, we open ourselves to novel biological insights that would remain difficult to identify without addressing the genome as a whole.

Computational tools are a powerful means to generate hypotheses that can be tested at the bench. Here, we sought to generate hypotheses regarding why identified missense mutations found to decrease complementation fitness are deleterious, looking to determine whether protein structure was impacted by the missense mutations using AlphaFold. While this tool was applied to a couple of examples in this work, AlphaFold’s utility should be investigated for larger datasets with known links between phenotype and observed missense mutations (importantly, one group has already tested this with green fluorescent protein variants linked to fluorescence data) (35). In this work, we used a graphic user interface (GUI) version of AlphaFold on Google CoLab. Use of GUI tools allows anyone to pick up the tool, regardless of their computational background. In this manner, we sought to use available computational tools in such a way as to make them most available to anyone seeking to use them for their own applications. While GUI tools are more generally usable, they often allow for fewer simultaneous tests and may not provide the user with the same depth of data output as the command-line version. Therefore, when a larger number of tests or additional data outputs are required, users should familiarize themselves with the command-line versions. While the examples used in this study with AlphaFold did not yield any testable hypotheses regarding missense mutations impacting protein structure, it is important not to overgeneralize protein structure to protein function. Here, a computational tool such as UniRep that is trained on related sequences and may be used to predict the likelihood of an amino acid occurring at a particular position in a sequence could prove to be more powerful (46). Further, as biology hypothesis generation increasingly relies on computational tools, it is important to recognize that, presently, much of our hypothesis testing must rely on benchwork (such as generating protein mutants and checking their structure through crystallography).

Improving protocols for recombineering BACs with ssDNA oligonucleotides can improve the speed and cost for creating targeted mutations within a piece of synthetic DNA for downstream applications in genome engineering work, critical as researchers look to expand the scope of projects being tackled (47, 48). Such mutations may include changes to genome segments to improve fitness, diversification to identify viable changes in a particular locus, and introduction of small genetic engineering target sites (such as PAM sites and restriction enzyme cut sites). Further, introduction of target mutations using MAGE makes such processes highly multiplexable. Such efforts also improve the accessibility of synthetic biology techniques for labs or research that may not be as well-funded, critical especially to early-stage principal investigators, or research into domains of biology less immediately useful or outside of geographic biology hotspots, therefore less likely to obtain substantial funding. As we move forward with large-scale genome engineering work, we must be cognizant that, at this time, the work we do is prohibitively expensive, in terms of cost, time, and labor. Just as we as synthetic biologists hold biocontainment as a characteristic of responsible research, we must consider frequent communication about challenges and possible solutions being faced in this research important as well (49). In this manner, we can present problems faced to the general scientific community, where diverse perspectives can provide unique, creative solutions. Further, transparency provides researchers looking to perform large-scale genome engineering with an understanding of challenges faced during the process, which in turn allows them to better prepare and to set realistic timelines for strain completion.

## DATA AVAILABILITY

AlphaFold2.1.0 is available on Google CoLab for academic use (https://colab.research.google.com/github/deepmind/alphafold/blob/main/notebooks/AlphaFold.ipynb). BioRender is available for academic use (https://biorender.com). Bowtie2 is available on the Web for academic use (https://github.com/BenLangmead/bowtie2). Geneious Prime 2022.0.2 is available for academic use (http://www.geneious.com/). PyMOL Version 2.4.2 is available for academic use (https://pymol.org/2/). RCSB Pairwise Structure Alignment is available on the Web for academic use (https://www.rcsb.org/alignment).

## SUPPLEMENTARY DATA

Supplementary Data are available with this article.

## ACKNOWLEDGEMENT

The authors wish to thank Kamesh Narasimhan, George Chao, and Chun-Ting Wu for helpful discussions regarding difficult target PCR amplification, statistical analysis techniques, and paper content, respectively.

